# Functional cardiac consequences of β-adrenergic stress-induced injury in the *mdx* mouse model of Duchenne muscular dystrophy

**DOI:** 10.1101/2024.04.15.589650

**Authors:** Conner C. Earl, Areli J. Javier, Alyssa M. Richards, Larry W. Markham, Craig J. Goergen, Steven S. Welc

**Affiliations:** Weldon School of Biomedical Engineering, Purdue University, West Lafayette IN, USA; Indiana University School of Medicine, IN, USA; Musculoskeletal Health Sciences Program, Indiana University School of Medicine, Indianapolis, IN USA; Division of Pediatric Cardiology, Riley Children’s Hospital at Indiana University Health, Indiana University School of Medicine, Indianapolis, IN; Department of Anatomy, Cell Biology & Physiology, Indiana University School of Medicine, Indianapolis IN, USA; Indiana Center for Musculoskeletal Health, Indianapolis IN, USA

**Keywords:** Duchenne muscular dystrophy, mdx, isoproterenol, 4DUS, cardiac strain, mouse model

## Abstract

Cardiomyopathy is the leading cause of death in Duchenne muscular dystrophy (DMD), however, in the *mdx* mouse model of DMD, the cardiac phenotype differs from that seen in DMD-associated cardiomyopathy. Although some have used pharmacologic stress to enhance the cardiac phenotype in the *mdx* model, many methods lead to high mortality, variable cardiac outcomes, and do not recapitulate the structural and functional cardiac changes seen in human disease. Here, we describe a simple and effective method to enhance the cardiac phenotype model in *mdx* mice using advanced 2D and 4D high-frequency ultrasound to monitor cardiac dysfunction progression *in vivo*. For our study, *mdx* and wild-type (WT) mice received daily low-dose (2 mg/kg/day) isoproterenol injections for 10 days. Histopathologic assessment showed that isoproterenol treatment increased myocyte injury, elevated serum cardiac troponin I levels, and enhanced fibrosis in *mdx* mice. Ultrasound revealed reduced ventricular function, decreased wall thickness, increased volumes, and diminished cardiac reserve in mdx mice compared to wild-type. Our findings highlight the utility of low-dose isoproterenol in *mdx* mice as a valuable model for exploring therapies targeting DMD-associated cardiac complications.

**SUMMARY STATEMENT:** This work introduces an improved method to model heart failure in mouse models of Duchenne muscular dystrophy and comprehensively describes underlying cellular and physiologic mechanisms using advanced imaging techniques.

## 1. INTRODUCTION

Duchenne muscular dystrophy (DMD) is a progressive X-linked skeletal and cardiac myopathy affecting 1 in 5000 live male births.(1) It is caused by mutation of the dystrophin gene resulting in the loss of functional protein.(2) Dystrophin is a large protein found at the inner surface of the sarcolemma linking myofibrillar actin to the cytosolic surface of muscle cell membranes providing structural support. The loss of dystrophin leads to striated muscle membrane weakness with increased susceptibility to damage caused by mechanical stress.(3) Myocyte damage can lead to cell death, chronic inflammation, and fibrotic replacement of contractile tissue further contributing to the destruction of muscles and magnitude of the disease.(4) Clinically, DMD patients present with skeletal muscle weakness and wasting, impaired motor function, loss of ambulation, respiratory insufficiency, and eventually pronounced cardiac disease.(5-7) Progress in treating respiratory insufficiency and complications related to the deterioration of skeletal muscles has extended lifespan leading to cardiomyopathy as the primary life-limiting factor affecting DMD patients.(8-10)

The *mdx* mouse, which carries a spontaneous mutation that results in the absence of dystrophin, is the most widely used pre-clinical animal model of DMD.(11, 12) Muscles from DMD patients and *mdx* mice feature asynchronous muscle damage, chronic inflammation, fibro-fatty degeneration, and diminished function.(12) However, these clinical symptoms are less severe in *mdx* mice than DMD patients exemplified by only a minor reduction in lifespan.(13) In particular, cardiac disease occurs especially late in the lifespan of *mdx* mice relative to human disease. Modest changes in left ventricular function frequently are not detected until 9-12 months of age in *mdx* mice, whereas nearly 60% of patients have some degree of left ventricular functional impairment by the age of 18.(14-17) A limitation of the *mdx* mouse is that even with advanced age they do not develop dilated cardiomyopathy as seen in DMD patients.(18) To advance the disease state, the genetic background of mice carrying the *mdx* mutation has been altered. The DBA/2J (D2)-*mdx* murine model demonstrates worsened skeletal and cardiac muscle pathologies.(19, 20) However, the DBA/2J mouse strain is susceptible to spontaneous cardiac calcinosis pathologies (21) and its suitability as a model for DMD is debated.(22, 23) Alternatively, double knockout mouse models have been generated that combine the dystrophin mutation of *mdx* mice with an additional gene mutation, such as the *mdx*-utrn double knockout mouse model, which also exhibit worsened cardiac pathologies.(24) Additional targeted mutations, however, can lead to a different molecular basis of disease from that which occurs in DMD patients.(18) With cardiomyopathy becoming a primary cause of morbidity and mortality in DMD patients, representative pre-clinical models are critically important for testing novel therapeutics and pathological mechanisms related to cardiac disease.(25)

Stressors that increase cardiac workload can induce cardiac myocyte injury and a more severe pathology in *mdx* mice. The β-adrenergic agonist isoproterenol has become a commonly used pharmacological stimulus of sarcolemmal injury in dystrophic cardiac myocytes.(26-29) Isoproterenol induces a sudden increase in heart rate and contractility stimulating mechanical stress resulting in prominent injury and eventual fibrotic replacement of myocardial tissue.(26-28) This pharmacological approach may be advantageous to other means of elevating cardiac work, such as exercise, due to the low capacity of *mdx* mice for physical activity and may better recapitulate human DMD without introducing additional confounding genetic variables.(30) However, isoproterenol dosing concentrations, routes of administration, and reported cardiac outcomes are highly variable and in many cases lead to early mortality. (28, 29, 31-35).

In this investigation, we perform a comprehensive evaluation of cardiac injury, remodeling, function, and their dynamics to isoproterenol challenge in wild-type and *mdx* mice. We test for these changes at 10-12 weeks of age, before the clinical onset of cardiac disease in *mdx* mice, focusing on adaptive changes to isoproterenol-induced injury before the development of verified pathologies.(14-16) We utilize 2-D and 4-D high-frequency ultrasound imaging to test relative changes in left ventricle morphology and function with isoproterenol challenge over time. We also report for the first time changes in β-adrenergic responsiveness by using ultrasound to measure acute changes in function before and after isoproterenol injection at baseline and after seven days of isoproterenol challenge. Additional experiments assay for changes in the molecular stress response, myocardial damage, and fibrosis with acute and chronic isoproterenol treatment. Overall, our findings support that dystrophin-deficient cardiac myocytes are acutely susceptible to isoproterenol injury resulting in distinct pathological and functional outcomes consistent with features of cardiac disease in DMD patients.

## 2. METHODS

### 2.1 Mice

All animal experiments were performed according to approved Institutional Animal Care and Use Committee protocols at Indiana University School of Medicine (IUSM) and Purdue University, conforming with federal ethical regulations and AALAC standards for animal testing and research. C57BL/10ScSn-Dmd^mdx^/J mice (*mdx*) and C57BL/10ScSn/J (wild-type) mice were purchased from The Jackson Laboratory (Bar Harbor, ME) and bred to maintain active colonies or directly used for experimentation.

### 2.2 Isoproterenol stimulation

Wild-type and *mdx* male mice, 10 to 12 weeks of age, received subcutaneous injections of isoproterenol (Sigma #I6504) dissolved in saline (2 mg/kg body weight). Mice undergoing acute isoproterenol stimulation received a single injection. Chronic isoproterenol stimulation included 10 consecutive daily injections of isoproterenol (Fig. 1A, B). A control group of mice were treated with an equal volume of saline to control for phenotypes associated with stress due to daily handling and injections.(36) Twenty-four hours after the final injection mice were sacrificed, hearts were dissected, and separated into apical and mid-chamber coronal sections divided using a stainless steel slicing matrix (Zivic). Sample sizes of n=4-5 per genotype for control, n=10 per genotype for acute isoproterenol, and n=5 per genotype for chronic isoproterenol were used per experimental group. For the imaging portion of the study, female wild-type (n=10) and *mdx* mice (n=10), 10-12 weeks of age, similarly received daily subcutaneous injections of 2 mg/kg/day isoproterenol (MedChemExpress #HY-B0468). Immediately after completing the baseline imaging and while the mice were still under isoflurane anesthesia, the first dose of isoproterenol was administered to each mouse to assess cardiac function and physiologic response to the stimulus exactly one-minute post-injection. Each mouse in these groups then received daily injections for 10 days, however the injection on day 7 was also performed during physiologic monitoring and imaging under anesthesia. After completion of the 10-day isoproterenol challenge, the mice were imaged one final time at day 14 of the study and subsequently sacrificed. At this time, we dissected the heart, then weighed and prepared the samples for histological analysis (Fig. 1C).

**Figure 1:**
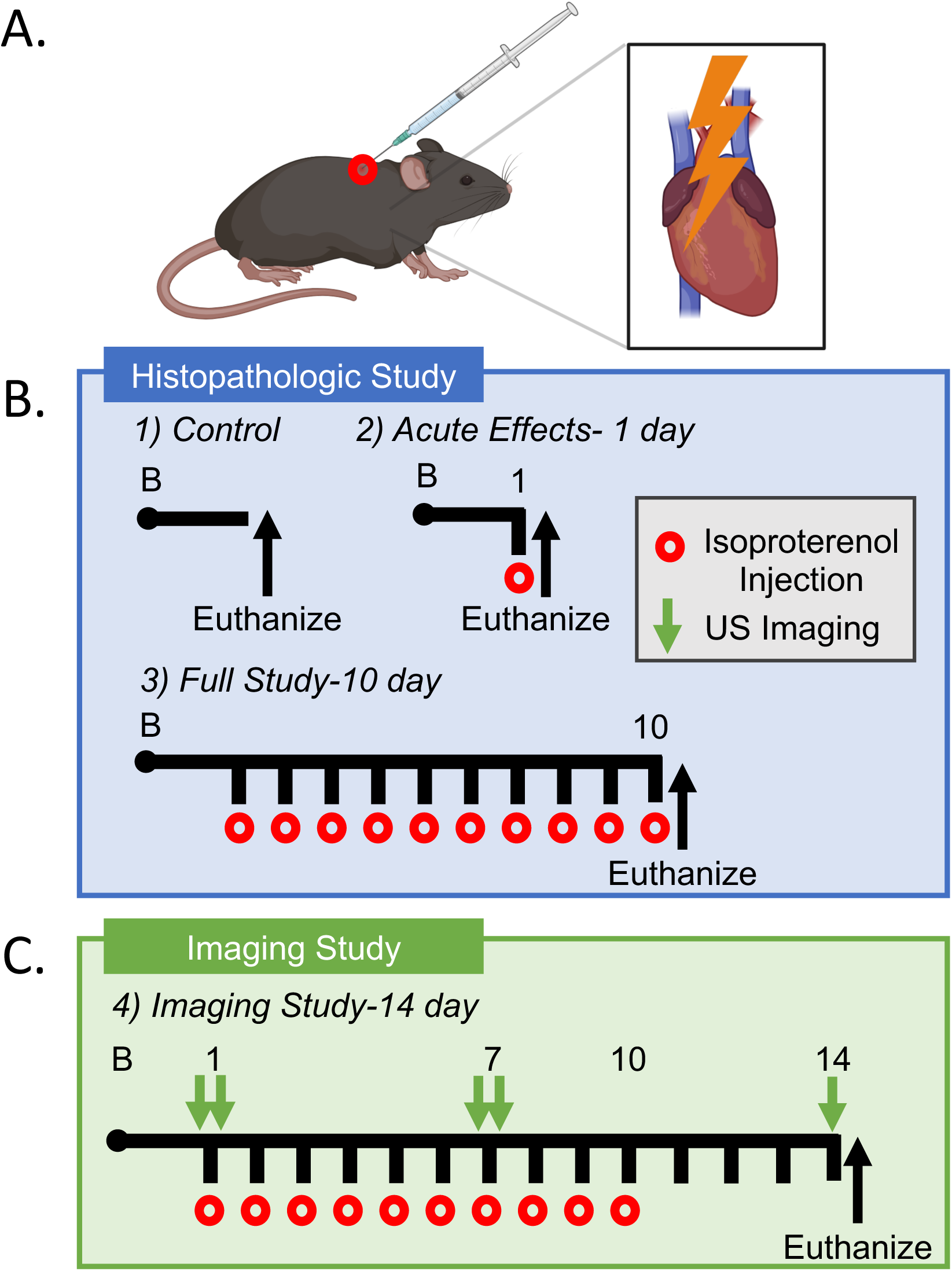
Study Overview. (A) Schematic representing subcutaneous injection of 2 mg/kg/day isoproterenol to induce β-adrenergic cardiac stimulation and injury. (B) Histopathologic study timeline. (C) Imaging study timeline.

### 2.3 Immunofluorescence

Mid-chamber ventricles were embedded in Optimal Cutting Temperature (OCT; Fisher Scientific), frozen in liquid nitrogen cooled isopentane, and cut to 10 µm thickness. To assess sarcolemmal damage in cardiac myocytes, coronal sections were fixed in ice cold acetone for 10 min. Sections were rinsed with PBS + 0.05% Tween20 (PBST) and blocked in 2% goat serum. Cardiac sections were then washed in PBST and labeled with FITC conjugated Wheat-Germ Agglutinin (1:500; Vector Labs #FL-1021-5) and Goat anti-Mouse IgM Alexa Fluor 594 (1:1000; Invitrogen #A21044) for 1 hour. Sections were mounted with Prolong Gold Antifade with DAPI (Invitrogen).

### 2.4 Histopathology

Fibrosis was assessed in myocardial sections by Sirius Red Fast Green (SRFG) stain, as previously reported with modification.^5^ Briefly, 10 µm thick mid-chamber coronal sections were fixed in ice cold acetone for 3 hours. Slides were rehydrated in 70% ethanol and rinsed with tap water. Sections were stained in 0.1% Direct Red 80 (Sigma #365548) and 0.1% Fast Green (Fisher #F88-10) in 1.3% Picric Acid (Sigma #P6744) for 25 min. and rinsed with tap water. Slides were sequentially dehydrated in 70% ethanol followed by 100% ethanol. Sections were cleared with Neo-Clear and mounted with NeoMount (Millipore). Fibrosis was confirmed with Masson’s Trichrome stains as previously described.(37)

### 2.5 Microscopy and Analysis

Imaging was performed in Zeiss ZenBlue 3.3 software and specimens were captured using on a Zeiss AxioObserver 7 inverted epifluorescence microscope with motorized stage, Zeiss AxioCam 105 color and Axiocam 506 monochromatic cameras. Whole tissue montages of WGA-IgM and SRFG were captured and stitched together with Zen Blue Autofocus and Tiles/Positions Modules. High magnification multi-color fluorescence images of WGA-IgM stained sections were captured as a Z stack of consecutive focal planes, deconvolved, and combined using the Zen Blue Z Stack and Extended Depth of Focus Modules. Collagen birefringence was assessed by transmitted light using a Polarizer D condenser fixed at a 90° angle and Analyzer Module Pol ACR P&C.

All images were deidentified and analyzed in a blinded fashion. Myocardial injury was quantified on sections stained with WGA and immunolabeled with fluorescence conjugated anti-IgM. The area of injury was determined by thresholding the IgM fluorescence positive area in ZenBlue 3.3 and expressing as a proportion of the total tissue area. To quantify fibrotic area, SRFG images were analyzed by measuring the Sirius Red positive pixel area using the color threshold function in ImageJ. Fibrotic area was then calculated as the proportion of Sirius Red positive area to total tissue area. Quantitative analysis of Masson’s Trichrome stain was also performed in ImageJ using the color segmentation plug-in to calculate the percentage of fibrosis relative to healthy tissue.

### 2.6 RNA isolation and quantitative PCR (QPCR)

The apex of the heart was homogenized in Trizol and RNA was extracted, separated with chloroform, and purified using RNeasy Plus kit with gDNA eliminator columns (Qiagen). RNA quantity and quality were measured using a microvolume NanoDrop One Spectrophotometer (Thermo Scientific). RNA was then electrophoresed on agarose gels and RNA quality was further assessed by 28S and 18S ribosome RNA integrity. RNA was reverse transcribed with Maxima H Minus cDNA Synthesis Master Mix (Thermo Scientific). Reaction and cDNA sample mixes were transferred to reaction plates by automation (Integra Assist Plus) and ran on a BioRad CFX384 QPCR system. QPCR experiments were designed as performed previously using established guidelines for experimental design, data normalization, and data analysis to maximize the rigor of quantifying the relative levels of mRNA.(38-40) The expression of each gene in wild-type control samples was set to one and other expression values were then scaled to that value. Primers are listed in Supplementary Table 1.

### 2.7 Serum cardiac troponin I (cTnI) ELISA

Blood was collected at the time of sacrifice and left to clot at room temperature for 30 min. Serum was separated by centrifugation and stored frozen at -20°C. Serum cardiac troponin I was measured using cTnI ELISA Kit (Life Diagnostics #CTNI-1-HS) according to the manufacturer’s instructions.

### 2.8 Ultrasound Imaging

Mice were anesthetized with 2% isoflurane in an induction chamber. Once anesthetized, we transferred each mouse to the supine position and secured it to a heated plate with electrodes (FUJIFILM VisualSonics, Toronto, CA) to monitor heart rate and respiratory rate. We delivered isoflurane through a nose cone maintained at 1.5% throughout the imaging procedure. Using a rectal temperature probe, we also maintained body temperature near 37±2°C.

Mice were imaged with the Vevo 3100 ultrasound system for small animals (FUJIFILM VisualSonics) using the MX550D linear array transducer (25-55 MHz). We used depilatory cream to remove abdominal and thoracic hair. We obtained transthoracic echocardiographic images of the left ventricle. The 2D images taken included a long- and short-axis ECG-gated kilohertz visualization (LAX EKV, SAX EKV), as well as long- and short-axis motion mode (LAX m-mode, SAX m-mode). A 4D image of the left ventricle was taken using a linear step motor by acquiring multiple SAX EKV images with a step size of 0.127 mm between images and a frame rate of 300 Hz. The lower bound of the image extended past the apical epicardium, and the upper bound was set to the ascending aorta.

On day 1 and day 7 of the isoproterenol injection protocol, we performed the full imaging protocol after which we administered the daily dose of subcutaneous isoproterenol (2 mg/kg) while the mouse was still under anesthesia. Approximately one minute following the injection, we obtained an LAX EKV image to compare left ventricular function changes prior to and immediately following isoproterenol challenge.

### 2.9 Statistical Analysis

All data are presented as mean±SEM. Statistical significance was calculated by unpaired *Student t*-test, one-way or two-way ANOVA with Tukey’s multiple comparison test to determine differences among multiple groups. Differences with a *P-*value < 0.05 were considered statistically significant.

## 3. RESULTS

### 3.1 Dystrophin-deficient cardiac myocytes are susceptible to acute sarcolemmal injury with isoproterenol treatment

The loss of dystrophin is associated with increased occurrences of cardiac myocyte damage, necrosis, and fibrotic replacement in patients with DMD. We examined cardiac myocyte damage in *mdx* mice at 10-12 weeks of age. Immunofluorescence analysis of mid-chamber myocardial cross-sections shows trace detection of dystrophic cardiac myocytes with IgM inclusion indicating that the occurrence of sarcolemmal injury does not differ from wild-type mice at this age (Fig. 2A). Next, wild-type and *mdx* mice were injected with a single dose of isoproterenol. Isoproterenol through its effects on rate and force of cardiac contraction, increases mechanical stress and is a well-accepted model of sarcolemmal injury in dystrophic mice.(26) Within 24-hours of acute isoproterenol (2 mg/kg) treatment approximately 8.5% of the myocardium was occupied by damaged IgM+ cardiac myocytes in *mdx* mice (Figs. 2A, B). We also tested the effects of chronic intermittent isoproterenol (2 mg/kg/day for 10 consecutive days) treatments on myocardial damage. Twenty-four hours after the final isoproterenol injection the area of the dystrophic myocardium occupied by IgM+ cardiac myocytes returned to control levels (Figs. 2A and Supplemental Figure 1A). In contrast, myocardial damage was negligible after acute and chronic isoproterenol treatments in wild-type mice (Fig. 2A, C and Supplemental Figure 1B). Importantly, survival rates were 100% in *mdx* and wild-type mice treated with isoproterenol (data not shown).

**Figure 2:**
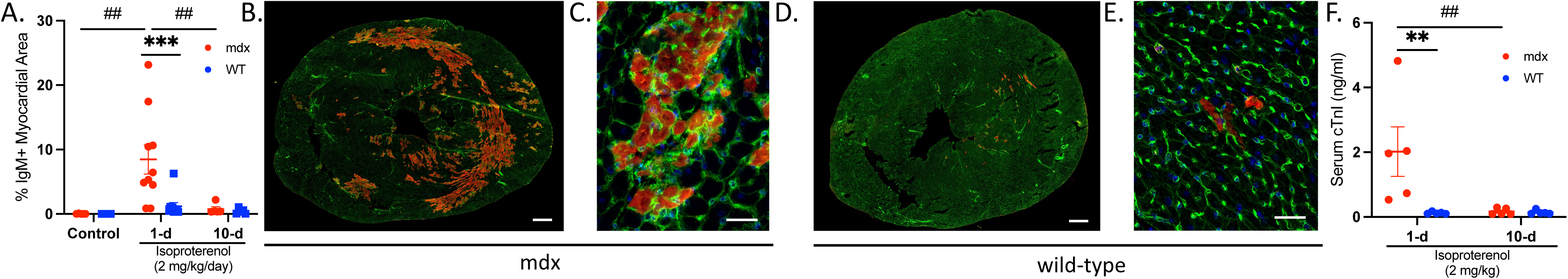
Isoproterenol promotes sarcolemmal injury in dystrophin-deficient cardiac myocytes. The acute and chronic effects of isoproterenol (2 mg/kg/day) were assessed in *mdx* and wild-type (WT) mice treated for 1- (n=10 per group) or 10 days (n=5 per group). Control mice were injected with an equal volume of saline for 10 days (n=5 *mdx* and n=4 WT). (A-D) To assess sarcolemmal damage, mid-chamber coronal sections of ventricles were immunolabeled with anti-IgM (red) and counterstained with wheat germ agglutinin (WGA; green) to visualize tissue morphology. (A) Cardiac injury was detected transiently in *mdx* mice after isoproterenol challenge with IgM+ cardiac myocytes occupying ∼8.5% of the myocardial area after a single isoproterenol treatment. Cardiac myocyte injury was scarcely detected in *mdx* mice after chronic isoproterenol administration or in WT mice across all conditions. (B-C) Representative *mdx* (B) and WT (C) whole ventricle cross-section and high magnification images show prominent areas of injury (red) after acute isoproterenol treatment in *mdx* mice. Bar=500 µm. (C) High-magnification images confirm that the IgM+ signal is localized to cardiac myocytes in *mdx* mice. (D) Serum cardiac troponin I (cTnI) levels were measured by ELISA as an independent assay of cardiac injury. Serum cTnI levels were also transiently elevated in *mdx* mice after acute isoproterenol treatment (n=5 per group). Data are presented as mean ± SEM. All *p* values are based on ANOVA with Tukey’s multiple comparison test. \*\**p*<0.01 and \*\*\**p*<0.001 versus WT within a treatment condition. ^##^ indicates *p*<0.01 between treatment groups within a genotype.

Finally, we measured serum cTnI concentrations with isoproterenol challenge. cTnI is almost exclusively expressed by cardiac myocytes and is a preferred biomarker for evaluation of myocardial injury in DMD.(41) Serum cTnI concentrations were greatly increased after acute isoproterenol stimulation in *mdx* mice relative to wild-type (Fig. 2D). However, serum cTnI levels were barely detectable in *mdx* mice after 10-days of isoproterenol stimulations. Serum cTnI concentrations did not change in wild-type mice with isoproterenol treatment. Consistent with our histological data, these findings suggest that 2 mg/kg/day of isoproterenol is insufficient to induce sarcolemmal damage in wild-type mice. Overall, our data support that dystrophic cardiac myocytes exhibit an acute increased susceptibility to damage with mechanical stress triggered by isoproterenol challenge.

### 3.2 Distinct cardiac growth and stress responses to isoproterenol challenge in dystrophin-deficient hearts

Because previous investigations have shown that isoproterenol stimulation induces cardiac stress and pathological growth (42) we compared its effects on indices of cardiac growth and stress in wild-type and *mdx* mice. We found that body weight was significantly higher in *mdx* (mean±SEM, 31.7±0.66g) compared to wild-type mice (26.3±0.61g), likely due to pseudo-hypertrophy of dystrophic skeletal muscles.(43, 44) Therefore, we used both heart weight normalized to tibia length (Fig. 3A) or body weight (Fig. 3B) as heart weight indices. Heart weight to tibia length ratio was higher in *mdx* mice compared to wild-type across all treatment conditions (Fig. 3A). Heart weight to tibia length (Fig. 3A) and heart weight to body weight (Fig. 3B) ratios increased 27% and 13% in wild-type mice with chronic isoproterenol stimulation relative to control, respectively. Whereas isoproterenol did not affect heart weight indices in *mdx* mice. The resulting isoproterenol stimulated increase in wild-type heart weight to body weight ratio exceeded that of *mdx* mice by 9%. These data indicate that at baseline some indexes of cardiac growth (heart weight to tibia length ratio) are increased in *mdx* mice compared to wild-type. However, isoproterenol stimulated increases in both heart weight indices (heart weight to tibia length or body weight ratios) in wild-type, but not *mdx* mice.

**Figure 3:**
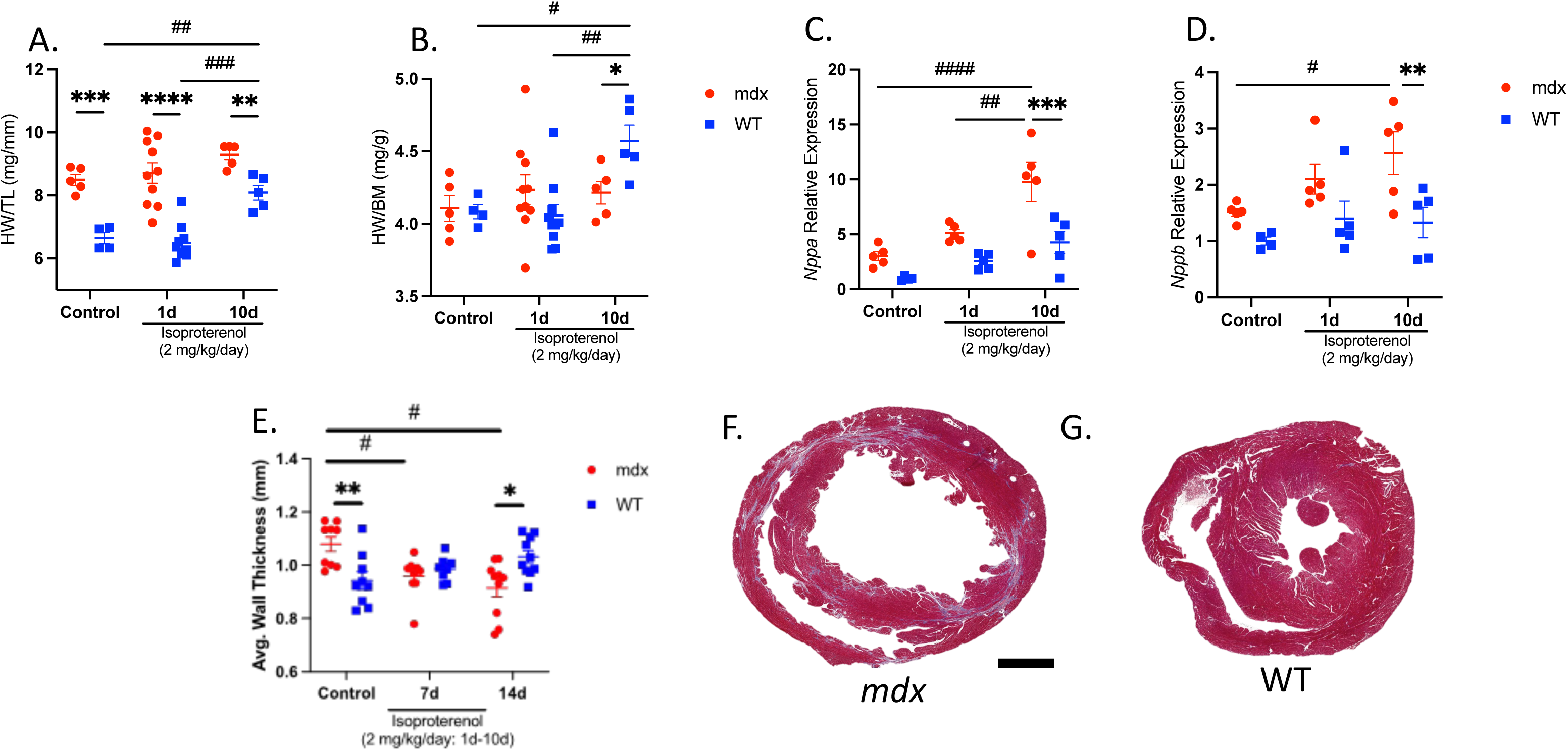
Distinct cardiac growth and stress responses to β-adrenergic stimulation in dystrophic hearts. Heart weight normalized to tibia length (HW/TL) and heart weight normalized to body mass (HW/BM) were used as indices of cardiac growth in *mdx* and WT mice treated for 1- (n=10 per group) or 10-days (n=5 per group), and control mice (n=5 *mdx* and n=4 WT). (A) HW/TL ratio was higher in *mdx* mice compared to WT in all conditions. Chronic isoproterenol treatment resulted in an increased HW/TL ratio in WT mice only. (B) HW/BM ratio increased with chronic isoproterenol treatment in WT mice relative to control and *mdx* mice. (C-D) QPCR analyses of *mdx* and WT ventricles expressed as mRNA levels relative to WT control. (C) *Nppa* and (B) *Nppb* mRNA expression increased in *mdx* mice with chronic isoproterenol challenge and relative to WT. (E) Ventricular wall thickness was measured at key time points by ultrasound (n=10 per genotype). At baseline, wall thickness was greater in *mdx* hearts relative to WT. However, in response to isoproterenol treatment wall thickness decreased in *mdx* mice becoming less thick than WT at day 14. Mid-chamber coronal sections of *mdx* (F) and WT (G) ventricles stained with Masson’s trichrome show characteristic fibrosis and thinning of the free wall in *mdx* mice at day 14. Bar=1 mm. Data are presented as mean±SEM. All *p* values are based on 2-way ANOVA with Tukey’s multiple comparison test. \**p*<0.05, \*\**p*<0.01, \*\*\**p*<0.001 and \*\*\**p*<0.0001 versus WT within a treatment condition. ^#^*p*<0.05, ^##^*p*<0.01, ^###^*p*<0.001 and ^####^*p*<0.0001 between treatment groups within a genotype.

Pathological cardiac growth and remodeling are also marked by increased expression of fetal genes *Nppa* and *Nppb* encoding atrial natriuretic peptide and brain natriuretic peptide, respectively.(45, 46) QPCR analysis was performed on ventricles from control and isoproterenol-treated wild-type and *mdx* mice. Chronic isoproterenol stimulation increased *Nppa* (Fig. 3C) and *Nppb* (Fig. 3D) mRNA expression by 3.3- and 1.7-fold in *mdx* mice compared to control, respectively. Consistent with the notion that isoproterenol-induced pathologies are exacerbated in dystrophin-deficient cardiac myocytes, *Nppa* and *Nppb* mRNA levels were 2.3- and 1.9-fold higher in *mdx* mice relative to wild-type with chronic isoproterenol treatment, respectively.

We also observed relative changes in myocardial wall thickness throughout the study. Using a 4-dimensional ultrasound (4DUS) analysis technique we characterized both the endocardial and epicardial surfaces using a 3-dimensional dynamic mesh over a characteristic cardiac cycle as described previously.(47, 48) We discovered an overall decrease in average myocardial thickness in *mdx* mice from baseline (1.08±0.08) to day 14 (0.92±0.10) following the 10-day course of isoproterenol treatments (*p=*0.029; Fig. 3E). In the wild-type group, we observed a slight, but not significant, increase in average myocardial thickness from baseline (0.94±0.10) to day 14 (1.03±0.07; *p=*0.088; Fig. 3E). This difference is also highlighted in the comparison between *mdx* and wild-type mice where *mdx* mice had a greater average myocardial thickness at baseline (*p*=0.006) but a reduced thickness at day 14 (*p*=0.010) after isoproterenol challenge (Fig. 3E). Reduced ventricular wall thickness was also apparent qualitatively on histologic examination at end-stage in *mdx* hearts (Fig. 3F,G; Supplemental Figure 2).

### 3.3 Isoproterenol promotes fibrotic replacement of the dystrophic myocardium

The increased occurrence of sarcolemmal damage with isoproterenol treatment in *mdx* mice could lead to cardiac myocyte cell death and fibrotic replacement of the myocardial tissue. We tested for changes in fibrosis by histology. The proportion of the myocardium occupied by collagen was increased by over 180% (Fig. 4A, B) in *mdx* mice with chronic isoproterenol treatment compared to control. No quantitative differences in fibrotic area were detected in wild-type mice consistent with our findings of negligible damage occurring in this group (Fig. 4A, C). Polarized light microscopy of myocardial areas positive for Sirius Red showed large birefringent areas indicating more densely bundled collagen strands in isoproterenol-treated *mdx* mice (Fig. 4D, F) relative to wild-type (Fig. 4E, G).

**Figure 4:**
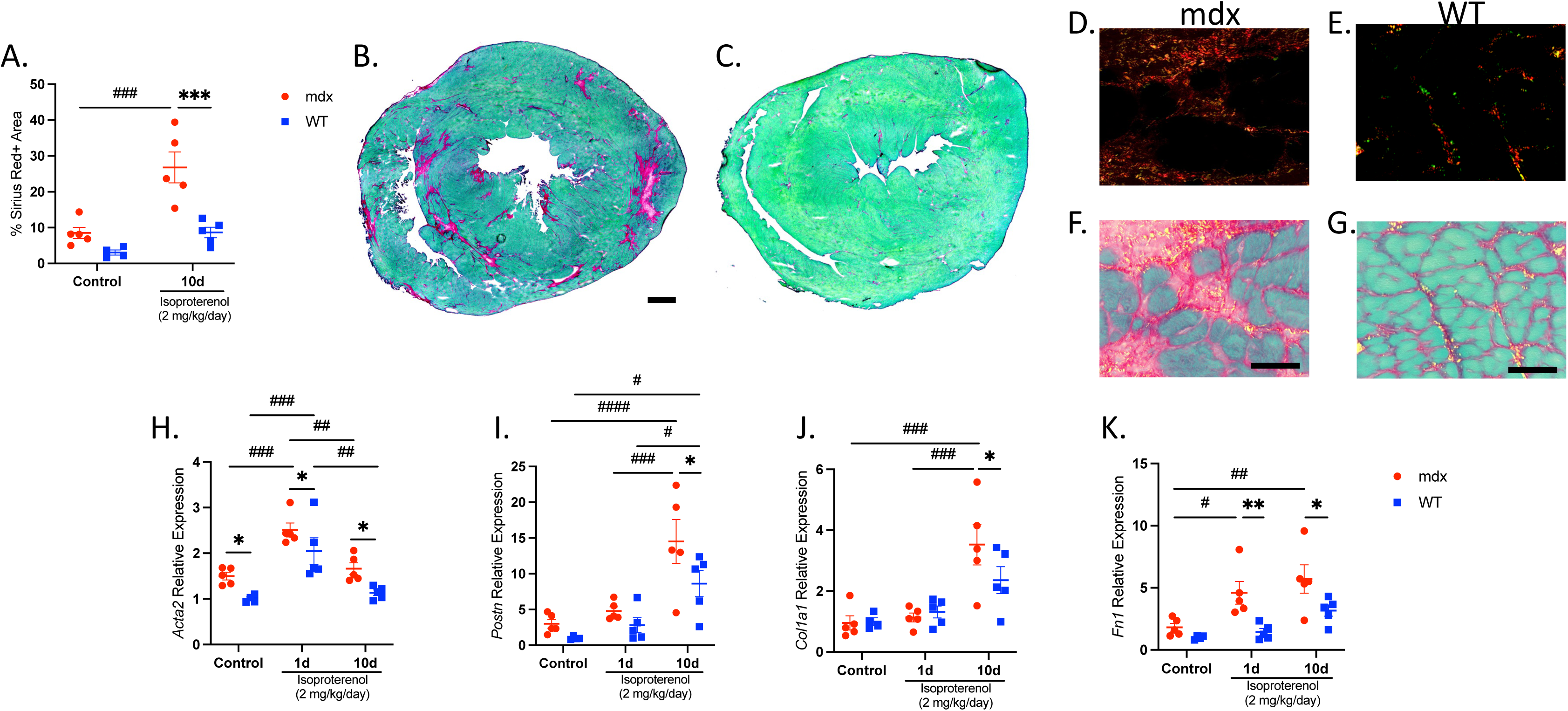
Isoproterenol promotes fibrotic replacement of the dystrophic myocardium. (A-G) Fibrotic area was measured on mid-chamber cross-sections stained with Sirius Red Fast Green stain (SRFG) in control (n=5 *mdx* and n=4 WT) and chronic isoproterenol treated (n=5 per genotype) mice. (A) The area of the myocardium occupied by collagen (red stain) is increased in (B) *mdx* mice after chronic isoproterenol challenge relative to control and (C) iso-treated WT mice. Bar=500 µm. (D-E) Collagen density was visualized by polarized light microscopy in iso-treated (D) *mdx* and (E) WT mice. (D-E) Fibrotic lesions of iso-treated *mdx* mice contained prominent birefringent areas indicating the presence of densely bundled collagen fibers (red and orange) in addition to the overall increase in collagen area. Representative overlay images of brightfield and polarized light (F) *mdx* and (G) WT ventricles. Areas of birefringence are pseudo-colored yellow to enhance image contrast in image overlays (F-G). (H-K) QPCR analyses of transcripts encoding markers of (H-I) activated fibroblasts and (J-K) connective tissue proteins. (H) *Acta2* mRNA levels were elevated in *mdx* mice relative to WT at baseline and with isoproterenol stimulation. (I) Induction of *Postn* transcripts was greater with isoproterenol stimulation in *mdx* relative to WT. Genes encoding extracellular matrix components (J) *Col1a1* and (K) *Fn1* were further increased in *mdx* mice relative to WT with iso-treatment. Data are expressed as mRNA levels relative to WT control group and presented as mean±SEM. All *p* values are based on ANOVA with Tukey’s multiple comparison test. \**p*<0.05, \*\**p*<0.01, and \*\*\**p*<0.001 versus WT within a treatment condition. ^#^*p*<0.05, ^##^*p*<0.01, ^###^*p*<0.001 and ^####^*p*<0.0001 between treatment groups within a genotype.

Next, we examined the expression of pro-fibrotic transcripts and their dynamics in response to isoproterenol challenge. QPCR analysis was performed to test for changes in transcripts encoding major extracellular matrix remodeling markers and connective tissue proteins. Alpha smooth muscle actin (*Acta2*) is a frequently used marker of cardiac myofibroblasts.(49) We observed a 1.6-, 1.2-, and 1.5-fold increase in *Acta2* mRNA expression in *mdx* mice compared to wild-type under control, acute and chronic isoproterenol stimulation conditions, respectively (Fig. 4H). *Acta2* mRNA expression levels were highest after acute isoproterenol stimulation increasing by 1.7- and 2-fold relative to control in *mdx* and wild-type mice, respectively. Periostin (*Postn*) is a matricellular protein also produced by activated cardiac fibroblasts whose activity is linked to fibrosis in muscular dystrophy.(50, 51) Chronic isoproterenol treatment induced a 4.9- and 8.6-fold increase in *Postn* gene expression in *mdx* and wild-type mice relative to control, respectively (Fig. 2I). However, *Postn* mRNA expression was 1.7-fold higher in the ventricles of *mdx* mice compared to wild-type after chronic isoproterenol stimulation. Collagen type 1 (*Col1a1*) and Fibronectin (*Fn1*) encode extracellular matrix components comprising fibrotic scar. *Col1a1* mRNA increased 3.5-fold in *mdx* mice with chronic isoproterenol treatment relative to control and the response was 1.5-fold higher than wild-type under identical conditions (Fig. 4J). In addition, *Fn1* mRNA expression increased 2.5- and 3.1-fold in *mdx* mice with acute and chronic isoproterenol treatment relative to control, respectively (Fig. 4K). The relative changes in *Fn1* transcripts were 3.2- and 1.8-fold higher in *mdx* mice compared to wild-type with acute and chronic isoproterenol treatment, respectively.

### 3.4 Left ventricular function and cardiac reserve decrease in mdx mice in response to isoproterenol challenge

To determine how isoproterenol injury affected left ventricular function *in vivo* we used both 2D and 4D high-frequency ultrasound imaging. Using 2D high-frequency ultrasound imaging we found a progressive decrease in left-ventricular ejection fraction (LVEF) in *mdx* mice from baseline (68±1%), day 7 (52±3%), and day 14 (41±3%), whereas we noted non-significant changes in wild-type mice during the same period with daily subcutaneous isoproterenol (2mg/kg/day) administration from day 1 to day 10 (Table 1; Fig. 5A). End-diastolic volume also increased in mdx mice from 53±2 µL at baseline to 86±9 µL at day 14 (*p*=0.01) as did end-systolic volume from 17±1 µL at baseline to 53±8 µL at day 14 (*p*<0.001). There were similarly no significant changes in end-diastolic or end-systolic volumes for the wild-type group over the course of the study (Fig. 5B,C). Volumetric changes at end-systole can also be qualitatively observed on imaging (Fig. 5F, representative images). There were no significant changes in cardiac output or heart rate over time for either group (Fig. 5D,E).

**Figure 5:**
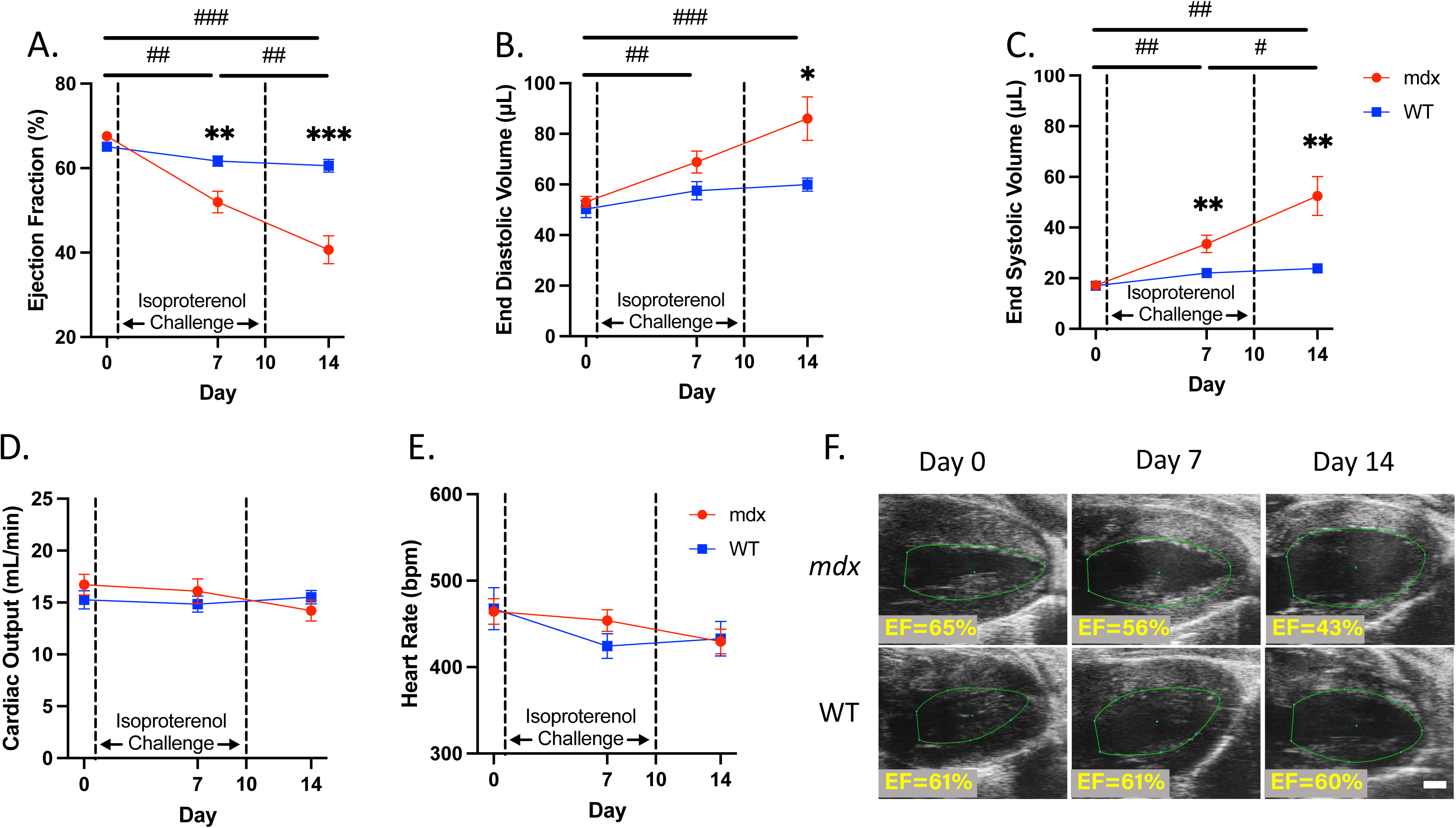
Cardiac function decreases with isoproterenol injury in dystrophic hearts. (A) LVEF in *mdx* mice was significantly reduced by day 7, and further reduced at day 14. (B) The left ventricular end-diastolic volume and (C) end-systolic volume in *mdx* mice were significantly greater than WT at day 14 signifying left ventricular dilation. (D) There were no significant changes in the cardiac output or (E) heart rate in either WT or *mdx* groups. (F) 2D ultrasound parasternal long-axis images at peak systole for both *mdx* and WT mice at day 0, day 7, and day 14 with left ventricular ejection fraction (EF) label. These findings suggest that the *mdx* mice exhibited dilated cardiomyopathy by day 14 when given an isoproterenol challenge. Data presented as mean±SEM. All *p* values are based on ANOVA with Tukey’s multiple comparison test. \**p*<0.05, \*\**p*<0.01, and \*\*\**p*<0.001 versus WT at a specified timepoint. ^#^*p*<0.05, ^##^*p*<0.01, and ^###^*p*<0.001 *mdx* difference between timepoints.

**Table 1:** Cardiac Function in WT and Dystrophic Mice with Isoproterenol Injury using 2D Ultrasound.

|  | Baseline |  |  | Day 7 |  |  | Day 14 |  |  |
| --- | --- | --- | --- | --- | --- | --- | --- | --- | --- |
|  | *mdx*<br>(n=10) | WT<br>(n=10) | *p* | *mdx*<br>(n=10) | WT<br>(n=10) | *p* | *mdx*<br>(n=10) | WT<br>(n=10) | *p* |
| LVEF (%) | 67.6±1.2 | 66.1±1.2 | 0.122 | 52.0±2.5 | 61.7±1.2 | 0.004* | 40.7±3.3 | 60.6±1.5 | <0.001* |
| EDV (μL) | 53.1±2.2 | 50.3±3.4 | 0.504 | 68.9±4.3 | 57.6±3.6 | 0.060 | 86.1±8.6 | 60.0±2.6 | 0.015* |
| ESV (μL) | 17.2±1.0 | 17.0±1.3 | 0.887 | 33.6±3.4 | 22.0±1.5 | 0.010* | 52.5±7.7 | 23.9±1.6 | 0.005* |
| CO (μL/min) | 16.7±1.0 | 15.3±0.9 | 0.295 | 16.1±1.2 | 14.8±0.8 | 0.400 | 14.2±1.0 | 15.5±0.6 | 0.298 |
| HR (beats/min) | 464±15 | 467±24 | 0.915 | 453±13 | 424±14 | 0.140 | 429±14 | 432±20 | 0.897 |

LVEF, left ventricular ejection fraction; EDV, left ventricular end-diastolic volume; ESV, left ventricular
end-systolic volume; CO, cardiac output; HR, heart rate

We also used ultrasound to observe the acute effects of isoproterenol injection for *mdx* and wild-type mice groups. At baseline, we observed and measured the cardiac response just prior to isoproterenol administration and approximately one minute following injection. While we observed a significant increase in absolute LVEF for both groups following isoproterenol injection at baseline (*p*<0.001), we found no differences in

ΔLVEF for both *mdx* (15.6±3.7%) and wild-type (15.4±5.6%) mouse groups (*p*=0.95) (Fig. 6 A,C). We also observed a robust heart rate increase immediately following isoproterenol administration compared to baseline for both *mdx* and wild-type mice with no significant difference in ΔHR between groups. However, at day 7 we did see a significant decrease in the ΔLVEF in the *mdx* group (7.3±4.0%) compared to the wild-type group (19.7±3.7%; *p*<0.001; Fig. 6 C,E; Supplemental Table 1). We also saw a significant decrease in ΔHR for the *mdx* group (*p*=0.01) compared to wild-type. Similarly, when comparing cardiac function metrics at baseline prior to and one minute following isoproterenol injection we saw no significant differences in ΔEDV, ΔESV, and ΔCO between *mdx* and wild-type groups (Supplemental Table 1). At day 7, there were differences between the *mdx* and wild-type groups for ΔESV (*p*=0.01), and ΔCO (*p*=0.03; Supplemental Table 2).

**Figure 6:**
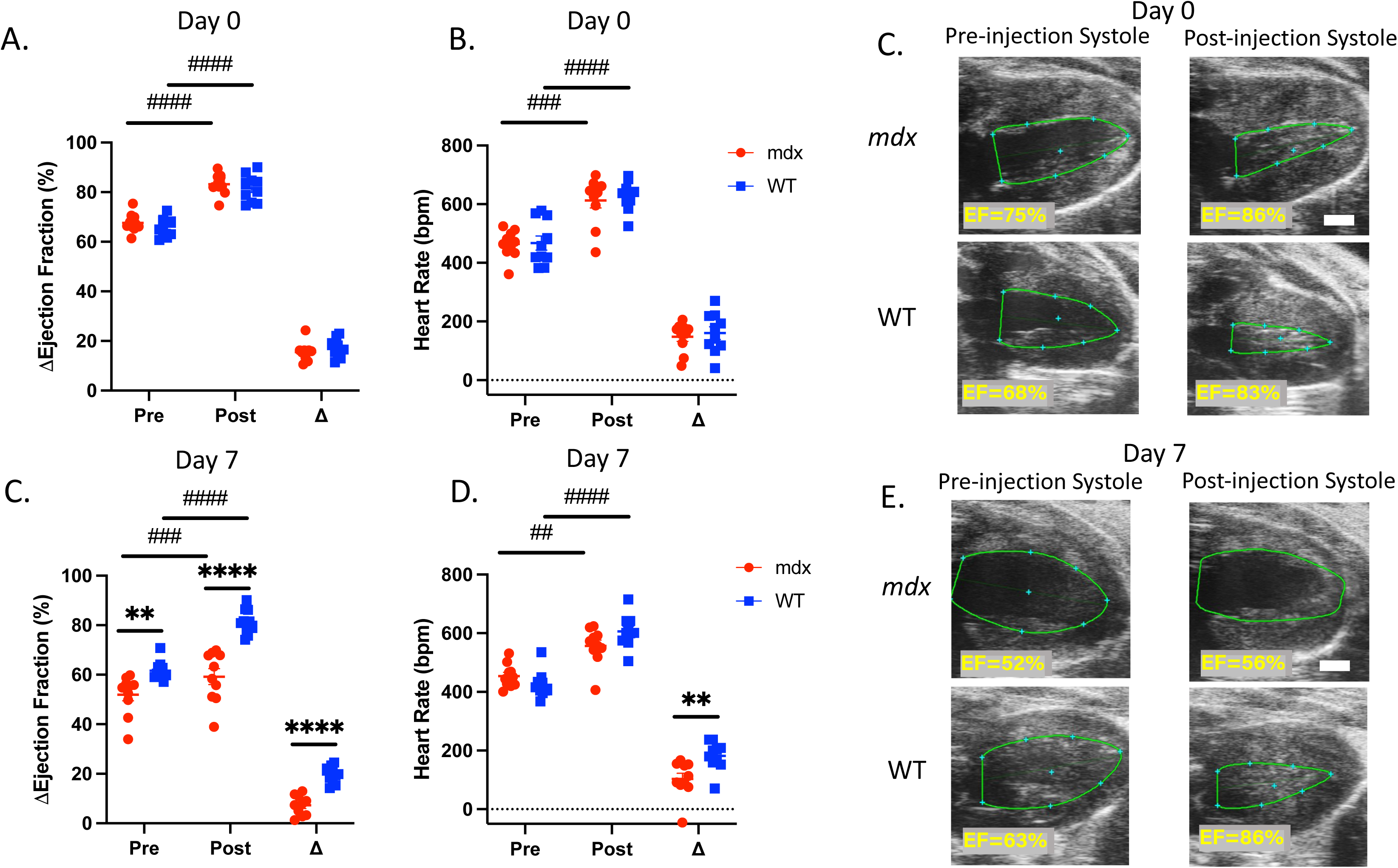
Cardiac response to isoproterenol challenge decreases over time in dystrophic hearts. (A,B) The initial compensatory response to an isoproterenol administration measured at exactly 1 minute before and 1 minute after the injection was apparent in the increase in heart rate and LVEF in both *mdx* and WT mice. (C) In the 2D ultrasound images of the left ventricle, this compensation can be seen as endocardial walls coming close together during systole. (C,D) After 7 days of exposure to isoproterenol injury, this compensatory response is impaired in the *mdx* mice, where the Δheart rate and ΔLVEF are reduced in comparison to WT. (E) There is little to no qualitative change in the left ventricular chamber in the *mdx* mice immediately after isoproterenol injection compared to WT. After prolonged exposure to an isoproterenol challenge, *mdx* mice’s ability to compensate for a single injection is impaired in comparison to the robust compensation of WT mice. Data presented as mean±SEM. All *p* values are based on ANOVA with Tukey’s multiple comparison test. \*\**p*<0.01, and \*\*\*\**p*<0.0001 versus WT. ^#^*p*<0.05, ^##^*p*<0.01, ^###^*p*<0.001, and ^####^*p*<0.0001 difference between pre and post injection.

**Figure 7:**
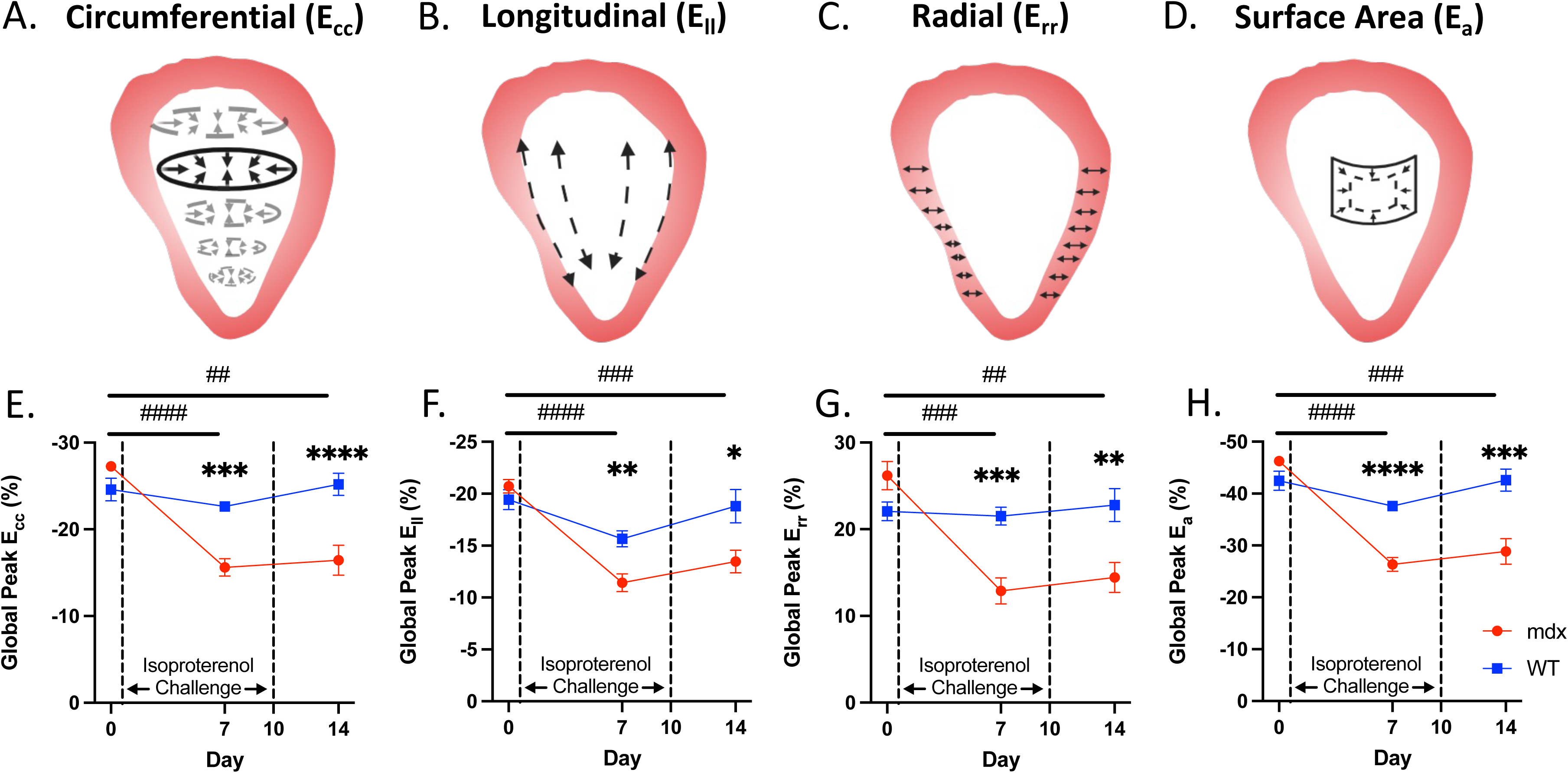
4D strain magnitude decreases in dystrophic hearts with isoproterenol injury. Global circumferential, longitudinal, radial, and surface area strain were calculated with 4DUS measurements (A-D). In the *mdx* mice with isoproterenol injury, the magnitude of strain was significantly lowered by day 7 (E-F). There was a slight recovery in ventricular strain by day 14, yet the strain magnitude remains significantly reduced. 4D strain decreases in dystrophic hearts when given an isoproterenol challenge. After the completion of the isoproterenol challenge, there is some functional recovery, but irreversible damage remains. Data presented as mean±SEM. All *p* values are based on ANOVA with Tukey’s multiple comparison test. \**p*<0.05, \*\**p*<0.01, \*\*\**p*<0.001 and \*\*\*\**p*<0.0001 versus WT at a specified timepoint. ^##^*p*<0.01, ^###^*p*<0.001, and ^####^*p*<0.0001 *mdx* difference between timepoints.

### 3.5 4D ultrasound myocardial strain magnitude decreases in dystrophic hearts with isoproterenol injury

Using our 4D strain technique, we estimated global circumferential (E_cc_), longitudinal (E_ll_), surface area (E_a_), and radial (E_rr_) strain components for *mdx* (n=10) and wild-type (n=10) mice at baseline, day 7, and day 14 after isoproterenol challenge on days 1-10. For example, in the *mdx* group, global average E_cc_ decreased from -27.3±1.0% at baseline to -15.6±3.1 (*p*<0.001) at day 7 and -16.4±5.4 (*p*=0.001) at day 14 while the wild-type had no significant changes from -24.6±4.1% at baseline to -22.6±1.3% (*p*=0.31) at day 7 and -25.2±4.0% (*p*=0.92) at day 14 (Fig. 6A). Global E_cc_ magnitude was also significantly decreased in *mdx* mice compared to wild-type at day 7 (*p*<0.001) and day 14 (*p*<0.001). We noticed a similar pattern of strain differences between *mdx* and wild-type groups for global E_ll_, E_a_, and E_rr_ (Fig. 6 B,C,D) as well as for regional strain patterns (Supplemental Table 3).

## 4. DISCUSSION

The results of our investigation demonstrate that mechanical stress induced by β-adrenergic stimulation advances disease state in *mdx* mice recapitulating cardiac pathologies present in DMD. Our findings show that before the acute onset of cardiac disease in *mdx* mice, cardiac myocytes are transiently susceptible to isoproterenol-induced injury, which is associated with increased fibrotic replacement of the myocardial tissue. The low-intensity isoproterenol stress protocol used here was insufficient to induce serum or histological markers of cardiac injury in normal hearts emphasizing the high sensitivity of dystrophin-deficient cardiac myocytes to mechanical stress. Additionally, the induction of transcripts encoding markers of cardiac stress, remodeling, and connective tissue proteins were potentiated in response to isoproterenol stimulation in dystrophic hearts. Distinctly, isoproterenol promoted thinning of the left ventricular wall and increased systolic and diastolic chamber volumes in *mdx* mice. Left ventricular ejection fraction progressively declined, while both cardiac reserve capacity and magnitude of myocardial strain also decreased in *mdx* mice in response to isoproterenol exposure. Finally, the isoproterenol challenge did not affect mortality and therefore dynamic and longitudinal changes in cardiac morphology and function are without survival bias. Collectively, these data support a reproducible model in which pharmacologically induced mechanical stress causes acute myocyte injury and progressive left ventricular dysfunction evolving into a form of dilated cardiomyopathy.

Relationships between isoproterenol-induced mechanical stress and dystrophic myocardial injury have been known for many years.(26) The magnitude of injury that we report, over 8% of the myocardium, is consistent with previous studies that also injected isoproterenol.(26, 29) Several investigations have shown that injections of higher concentrations of isoproterenol can result in a more severe injury that is also associated with a higher incidence of mortality in *mdx* mice.(52-54) Damaged myocytes are cleared from the dystrophic heart within one week of a single isoproterenol injection.(27) Our observation that cardiac injury was negligible in *mdx* mice after ten daily isoproterenol treatments suggests that its effects on dystrophic myocyte damage are transient. In support of this observation, low incidences of cardiac injury were also reported in *mdx* mice implanted with osmotic pumps for 14 or 28 days of continuous isoproterenol infusion.(28) However, we also find that the expression of cardiac stress genes, *Nppa* and *Nppb,* are highest in *mdx* hearts after chronic isoproterenol treatment. That observation could suggest that dystrophic hearts are in a state of elevated stress despite only an acute susceptibility to sarcolemmal injury with sustained isoproterenol treatment.

Our findings also indicate gross differences in left ventricular remodeling and morphology in dystrophin-deficient hearts. We observed a higher ratio of heart weight to tibial length in *mdx* mice relative to wild-type. However, this ratio only increased significantly in wild-type mice in response to isoproterenol treatment, while it remained relatively unchanged in the *mdx* group. In our 4DUS studies, we observed that global average systolic wall thickness was initially higher in *mdx* mice relative to wild-type. Still, isoproterenol treatment led to significant thinning of the ventricular wall in *mdx* mice, whereas there was an insignificant increase in wall thickness in wild-type mice. We also noted decreased left ventricular wall thickness in *mdx* mice in our histological studies. Furthermore, we found that isoproterenol stimulated an increase in end-diastolic and end-systolic left ventricular chamber volumes in *mdx* mice. Due to reported lethality,(28, 32) the isoproterenol dose (2 mg/kg/day) used in the present study is considerably lower than standardized doses (30 mg/kg/day) known to induce hypertrophy in normal hearts and likely explains why we observe only a mild hypertrophic phenotype in wild-type mice.(55-58) Collectively, these data support that the *mdx* model’s susceptibility to isoproterenol injury and fibrotic replacement of viable cardiac tissue leads to ventricular dilation.

To our knowledge, this is the first study to use 4D high-frequency ultrasound to evaluate progressive cardiac functional, morphological, and kinematic changes due to mechanical stress-induced injury in dystrophin-deficient hearts *in vivo*. Consistent with previous reports(14-16, 59, 60), our findings show no significant differences in baseline cardiac function in 10-12 week-old *mdx* mice. However, after initiating isoproterenol treatment, left ventricular ejection fraction progressively decreased in *mdx* mice. Despite these changes, we found that cardiac output is preserved throughout the study, which may indicate that the functional changes we observe are enough to compensate for metabolic demand. When this demand is challenged acutely with isoproterenol, we begin to see evidence of decompensation as both ΔHR and ΔCO are significantly decreased in the *mdx* group after 7-days of repeated isoproterenol doses. These results appear consistent with previous work showing that reduced acute cardiac β-adrenergic response is an early indicator of heart disease at the compensatory stage.(59) This reduction in cardiac reserve capacity may occur due to the loss of myocytes with the progressive onset of heart disease or in response to isoproterenol injury in *mdx* mice. We also noted a decrease in 4DUS strain magnitude for global circumferential, longitudinal, radial, and surface area strain components in dystrophic hearts in every region examined as well as globally suggesting that the response to isoproterenol injury affected the entire myocardium. In our histology studies, we observed characteristic diffuse fibrotic lesions throughout the myocardium potentially accounting for changes in myocardial kinematics measured with strain. Each strain component was reduced at day 7 relative to baseline, but there was not a further reduction from day 7 to day 14, as seen with the ejection fraction. One possible explanation for this is that our strain metrics may be able to indicate the full extent of myocardial damage earlier than other global or traditional metrics. Alternatively, this finding could indicate a partial recovery of cardiac contractility at day 14 following the termination of isoproterenol treatments at day 10. Future work will be needed to elucidate these possibilities.

While others have used isoproterenol to progress cardiac disease state in *mdx* mice. (28, 29, 31-35), the reported cardiac functional outcomes of isoproterenol injury are highly variable from study to study, in part due to differences in cardiac readouts, as well as unique isoproterenol protocols differing in dosage, frequency, route of administration, and method of delivery affecting comparisons between studies. For example, three sequential dosages of isoproterenol (0.35 mg/kg) over 12 hours caused acute systolic and diastolic hemodynamic dysfunction.(29) Echocardiographic assessment showed a modest 9% reduction in ejection fraction and a 13% decrease in shortening fraction in *mdx* mice 5 days after completing 5 daily doses of isoproterenol (3 mg/kg/day).(31) A decrease in ejection fraction and shortening fraction was seen in *mdx* mice one day after implantation of an osmotic pump for continuous delivery of isoproterenol (4 mg/kg/day), however high mortality rates prevented further characterization.(32) Osmotic pump delivery of a high concentration of isoproterenol (30 mg/kg/day) for 7 or 14 days did not affect ejection fraction or shortening fraction in *mdx* mice.(33) In contrast, another report found that *mdx* mice succumbed within 24 hours of implantation of pumps delivering 1, 5, 10, 15, and 30 mg/kg/day, but were able to mostly tolerate 0.5 mg/kg/day. Two weeks of delivery at this low dose stimulated an increase in the shortening fraction, however, these changes did not persist returning to control levels after 4 weeks.(28) Thus, systolic dysfunction is not observed in *mdx* mice receiving non-lethal doses of isoproterenol via an osmotic pump, indicating that the mode of delivery may account for some of the variation in functional outcomes between studies. However, the findings in our study are in general agreement with others showing that the injection of isoproterenol promotes systolic dysfunction in *mdx* hearts.

The mdx model of human DMD has been extensively phenotyped and the limitations of the model, as it relates to cardiomyopathy, have been described. Numerous studies have proposed therapies to ameliorate disease in the mdx mouse, even reporting that the mouse has been cured.(61-64) However, these therapies have not been translated to human clinical trials, especially as it relates to cardiomyopathy. Current guidelines suggest a proactive therapeutic approach of therapies largely derived from evidence-based in large-scale adult cardiomyopathy and heart failure trials.(65, 66) There is a need for therapies specific to the loss of dystrophin and the subsequent pathophysiologic cascade that may differ from cardiomyopathy of other etiologies.(67, 68) While no murine model can perfectly recapitulate the cardiomyopathy progression in DMD(69, 70), this model of β-adrenergic stress-induced injury has benefits that could lead to the rapid discovery and translation of DMD cardiomyopathy-specific therapies to bring to human clinical trials.

## 5. CONCLUSIONS

This study reports that our improved method of low-dose β-adrenergic stimulation is pathogenic for the acceleration of cardiac dysfunction in *mdx* mice. The primary, novel finding of this work is that repetitive isoproterenol injections cause a progressive reduction in ejection fraction, reduced acute cardiac β-adrenergic responses, and reductions in cardiac strain magnitude, both regionally and globally without causing mortality in our model. Our study also shows that isoproterenol reduced ventricular wall thickness and increased chamber diameter in *mdx* hearts. This is functionally important in the context of DMD in which heart disease progresses to a form of dilated cardiomyopathy. These data highlight the cardiac functional consequences of the acute vulnerability of dystrophin-deficient cardiac myocytes to mechanical stress. An additional strength of this work is the establishment of an experimental platform with a simple intervention, daily subcutaneous isoproterenol injections, and well-defined functional outcomes that could be used to evaluate therapies targeting contraction-induced damage or associated responses in dystrophic hearts.

## Declaration of Interest

CJG is a paid consultant of FUJIFILM VisualSonics Inc.

## Funding

Research reported in this publication was supported by the National Heart Lung and Blood Institute under award numbers F30HL162452 (to CCE) and R01HL158647 (to SSW) and the National Institute of Arthritis Musculoskeletal and Skin Disease T32AR065971 (to AJJ) of the National Institutes of Health. Additional support was provided by the Indiana University School of Medicine and Purdue University College of Engineering in Medicine Pilot Project Initiative (to CGJ and SSW) and the Riley Children’s Foundation (to CJG and LWM). The content is solely the responsibility of the authors and does not necessarily reflect the official views of the National Institutes of Health.

## Declaration of generative AI and AI-assisted technologies in the writing process

During the preparation of this work, the authors used ChatGPT(OpenAI, San Francisco, California) in some minor instances to rephrase and/or summarize text in order to improve readability. After using this tool, the authors reviewed and edited the content as needed and take full responsibility for the content of the publication.

## Author Contributions

Conceptualization and experimental design (CCE, AJJ, LWM, CJG, SSW); methodology and performance of experiments (CCE, AJJ, AMR), data analysis (CCE, AJJ, AMR, LWM, CJG, SSW); supervision (CJG, SSW); writing of the manuscript (CCE, AJJ, AMR, LWM, CJG, SSW). All authors have read and agreed to the published version of the manuscript.

## Data availability

The data presented in this work is freely available upon reasonable request.

## SUPPLEMENTAL

**Supplemental Table 1:**
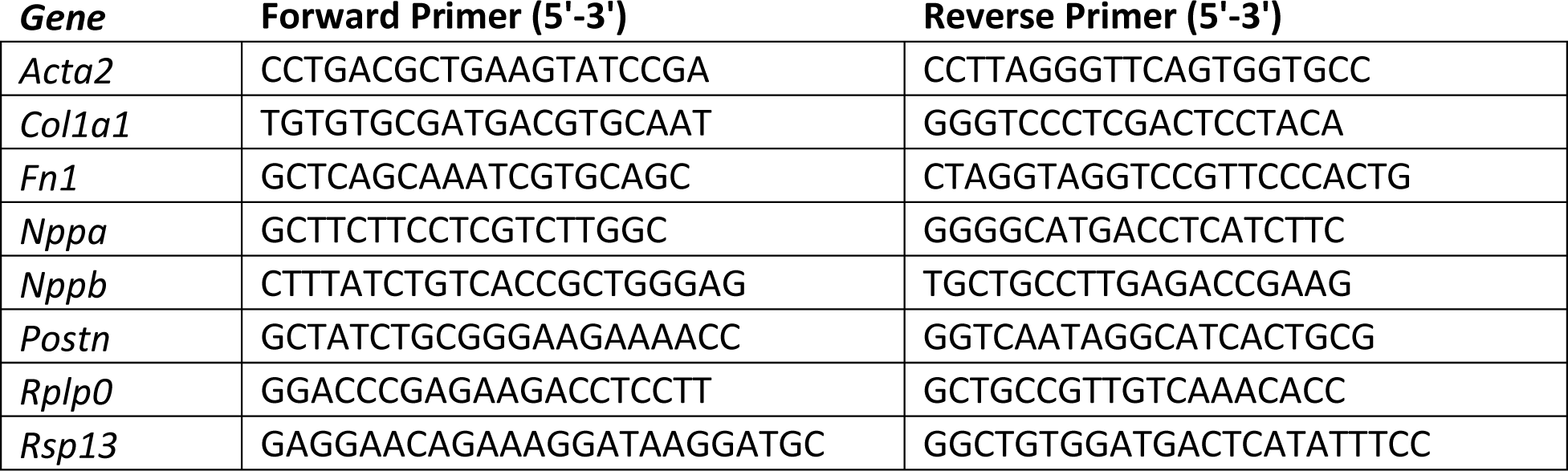
qPCR Primers.

**Supplemental Figure 1:**
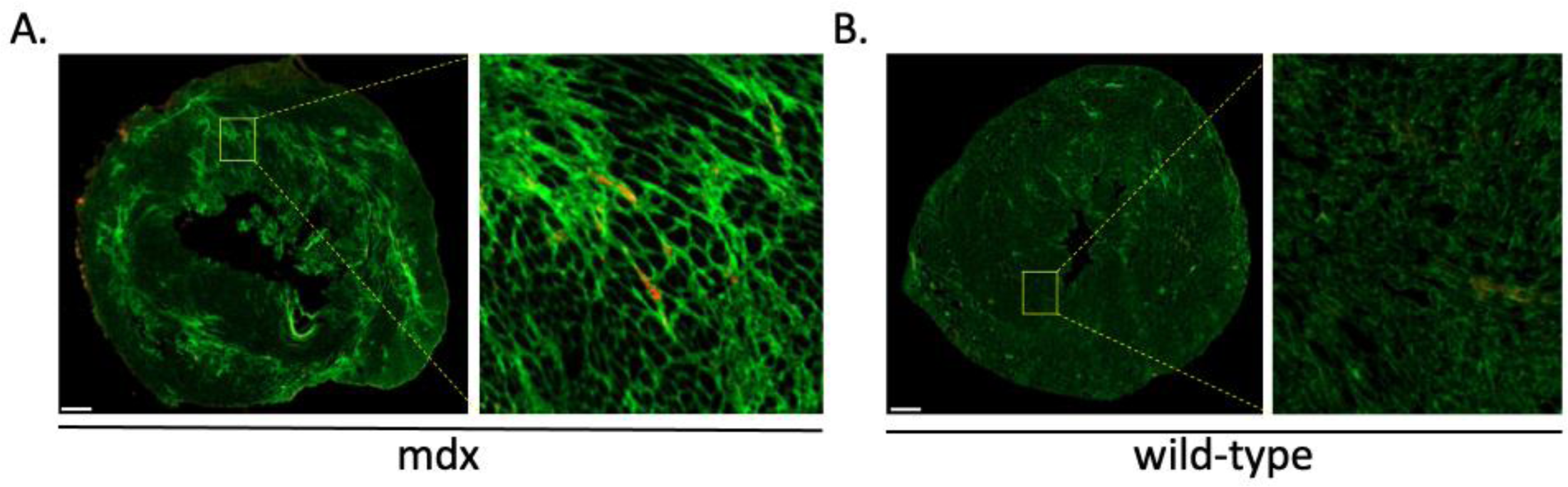
Cardiac injury is not sustained with chronic isoproterenol treatment. Representative images of *mdx* (A) and WT (B) myocardial cross-sections with chronic isoproterenol treatment immunolabeled with anti-IgM (red) and counterstained with WGA (green). Whole ventricle montages show trace IgM signal in (A) *mdx* and (B) WT ventricles. Bar=500 µm. High magnification images show rare IgM positive signal in (A) *mdx* and (B) WT ventricles. Bar=20 µm.

**Supplemental Figure 2:**
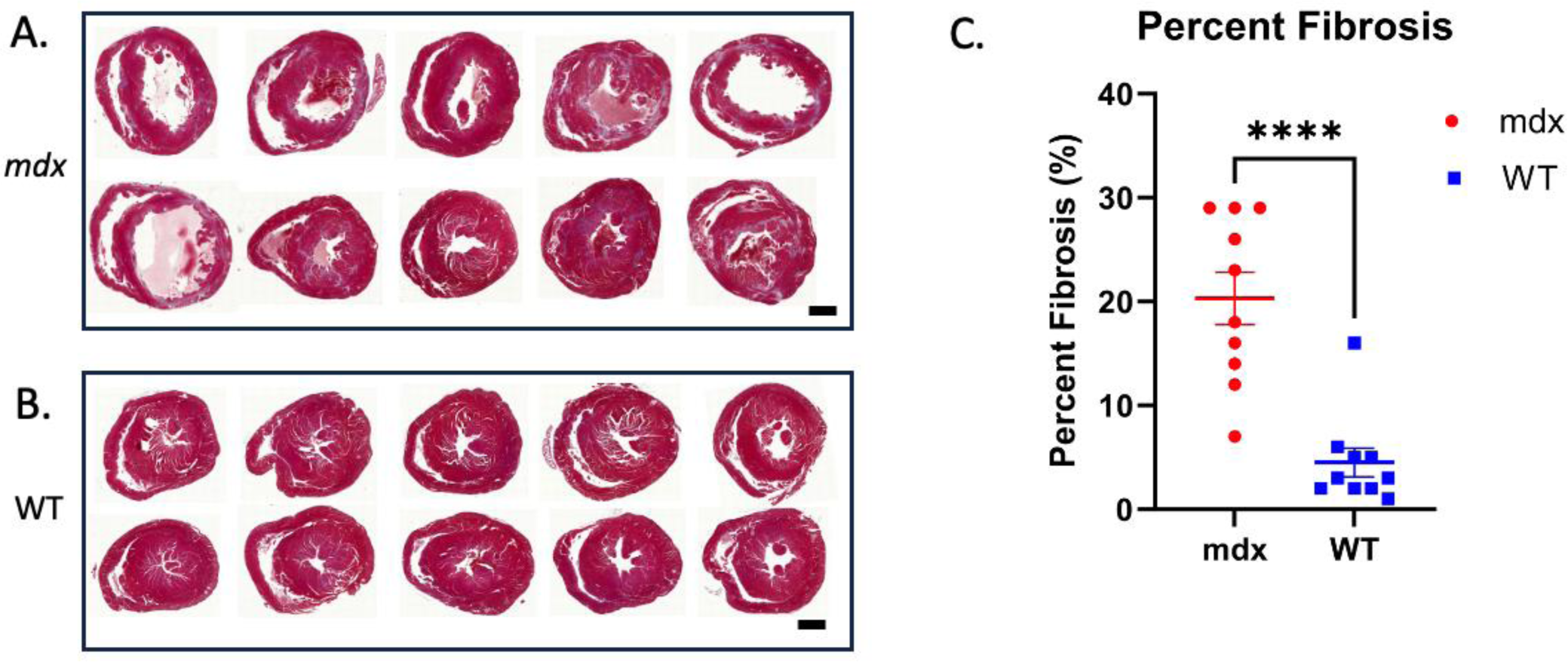
Fibrosis burden increased in *mdx* mice compared to wild-type with chronic isoproterenol treatment. (A) Masson’s trichrome stained mid-ventricle cross sections from *mdx* mice and (B) wild-type mice exposed to chronic isoproterenol treatment. (C) Percent fibrosis differences between groups. \**p*<0.05, Bar=1 mm.

**Supplemental Table 2:** Isoproterenol challenge response in *mdx* and wild-type mice with chronic isoproterenol treatment using 2D ultrasound.

|  | Baseline |  |  | Day 7 |  |  |
| --- | --- | --- | --- | --- | --- | --- |
|  | mdx<br>(n=10) | WT<br>(n=10) | *p* | mdx (n=10) | WT (n=10) | *p* |
| ΔLVEF (%) | 15.6±1.2 | 15.4±1.8 | 0.944 | 7.3±1.3 | 19.7±1.1 | <0.001* |
| ΔEDV (μL) | -20.5±2.7 | -16.5±4.0 | 0.376 | -11.1±2.9 | -16.5±2.8 | 0.230 |
| ΔESV (μL) | -11.6±1.0 | -10.6±1.6 | 0.611 | -8.6±1.3 | -14.4±1.4 | 0.005* |
| ΔCO (μL/min) | -0.2±1.0 | 1.9±0.9 | 0.118 | 2.1±1.0 | 5.2±1.0 | 0.023* |
| ΔHR (beats/min) | 148±16 | 161±22 | 0.641 | 103±20 | 182±15 | 0.004* |

**Supplemental Table 3:** Regional 4DUS strain comparison between wild-type and *mdx* mice.

|  |  | Baseline |  |  | Day 7 |  |  | Day 14 |  |  |
| --- | --- | --- | --- | --- | --- | --- | --- | --- | --- | --- |
|  |  | mdx<br>(n=10) | WT<br>(n=10) | *p* | mdx<br>(n=10) | WT<br>(n=10) | *p* | mdx<br>(n=10) | WT<br>(n=10) | *p* |
| Circumferential<br>($E_{cc}$) | Basal | -25.1±0.8 | -22.8±1.5 | 0.223 | -14.4±1.2 | -20.9±0.9 | <0.001* | -16.4±1.6 | -24.0±1.4 | 0.003* |
|  | Mid | -27.4±0.8 | -25.0±1.1 | 0.115 | -15.4±1.1 | -23.6±0.4 | <0.001* | -15.7±1.9 | -26.2±1.1 | <0.001* |
|  | Apical | -27.0±1.1 | -25.9±1.3 | 0.525 | -17.1±0.8 | -23.4±0.8 | <0.001* | -17.2±1.5 | -25.3±1.4 | 0.002* |
|  | Global | -26.5±0.7 | -24.6±1.2 | 0.213 | -15.6±0.9 | -22.6±0.4 | <0.001* | -16.4±1.6 | -25.2±1.2 | <0.001* |
| Longitudinal<br>($E_{ll}$) | AFW | -20.2±0.8 | -19.5±0.8 | 0.590 | -12.2±0.9 | -16.4±0.6 | 0.002* | -14.0±1.1 | -19.9±1.6 | 0.011* |
|  | A | -19.0±0.5 | -18.4±0.9 | 0.534 | -10.4±0.8 | -14.7±0.9 | 0.002* | -12.4±1.2 | -17.6±1.6 | 0.022* |
|  | AS | -18.8±0.6 | -17.9±0.9 | 0.414 | -10.2±0.8 | -14.3±0.8 | 0.003* | -12.8±1.0 | -16.5±1.5 | 0.074 |
|  | PS | -19.3±0.7 | -18.8±1.2 | 0.727 | -10.3±0.7 | -13.9±0.8 | 0.004* | -12.6±1.0 | -16.9±1.3 | 0.028* |
|  | P | -21.3±0.9 | -20.8±1.0 | 0.756 | -12.1±0.8 | -16.9±0.8 | <0.001* | -14.1±1.1 | -20.8±1.5 | 0.004* |
|  | PFW | -21.2±0.9 | -21.2±0.9 | 0.975 | -13.4±1.0 | -17.7±0.8 | 0.006* | -14.9±1.0 | -21.3±1.5 | 0.005* |
|  | Global | -20.0±0.7 | -19.4±0.9 | 0.080 | -11.4±0.8 | -15.7±0.7 | 0.002* | -13.5±1.0 | -18.8±1.5 | 0.013* |
| Radial<br>($E_{rr}$) | Basal | 14.9±1.4 | 11.1±0.5 | 0.030* | 7.3±0.7 | 9.6±0.7 | 0.034* | 7.5±1.1 | 10.7±1.1 | 0.076 |
|  | Mid | 32.8±2.2 | 27.2±1.5 | 0.064 | 15.8±2.0 | 27.8±1.4 | <0.001* | 16.9±2.8 | 28.8±2.3 | 0.006* |
|  | Apical | 32.7±2.0 | 30.7±2.4 | 0.562 | 17.0±2.2 | 30.0±1.9 | <0.001* | 21.1±1.9 | 32.1±3.2 | 0.014* |
|  | Global | 26.0±1.6 | 21.7±1.0 | 0.053 | 12.7±1.4 | 21.2±1.0 | <0.001* | 14.2±1.7 | 22.5±1.8 | 0.005* |
| Surface Area<br>($E_a$) | Basal | -46.7±1.5 | -44.9±1.9 | 0.472 | -28.5±1.1 | -39.9±1.4 | <0.001* | -30.5±2.2 | -43.8±2.0 | <0.001* |
|  | Mid | -44.7±1.3 | -41.5±1.6 | 0.167 | -25.3±1.3 | -37.0±0.9 | <0.001* | -27.6±2.5 | -42.3±2.0 | <0.001* |
|  | Apical | -43.3±1.3 | -40.3±1.8 | 0.219 | -24.7±1.5 | -35.0±1.0 | <0.001* | -28.3±2.3 | -41.2±2.3 | 0.002* |
|  | Global | -45.2±1.2 | -42.5±1.7 | 0.252 | -26.4±1.2 | -37.7±0.9 | <0.001* | -28.9±2.3 | -42.6±2.0 | <0.001* |

AFW, anterior free-wall; A, anterior; AS, anterior septal; PS, posterior septal; P, posterior; PFW, posterior free-wall

## Notes

### Competing Interest Statement

Craig J. Goergen is a paid consultant of FUJIFILM VisualSonics Inc.

